# MARiO: predicting cancer variant pathogenicity by integrating *in silico* evaluation and patient-level mutational contexts

**DOI:** 10.64898/2026.07.07.736919

**Authors:** Haruko Nakagawa, Takashi Kamatani, Naoya Ishibashi, Satoru Aoyama, Michina Morioka, Fuyuki Miya, Sadakatsu Ikeda

## Abstract

Comprehensive genomic profiling (CGP) supports precision medicine in cancer care, but accurate assessment of missense variant pathogenicity, especially for variants without established consensus, remains challenging. Various computational tools have been developed for variant functional prediction, but most current tools rely solely on variant-level features and do not capture the clinical context of individual patients. To address this limitation, we developed MARiO (Missense Alteration Risk for Oncogenicity), a machine-learning model that integrates variant-level features and patient-level clinical and genomic contexts to effectively predict the pathogenicity of missense variants in cancer. We collected a total of 10,642 missense variants from 1271 patients, and evaluated candidate features for their association with variant pathogenicity, identifying informative features including *in silico* functional predictions, population allele frequency, variant allele frequency, and tumor mutational burden. Using these selected features, MARiO was developed with extreme gradient boosting. The model integrates multiple *in silico* prediction tools and patient-specific genomic contexts while accommodating missing values frequently observed in real-world CGP datasets. MARiO outperformed existing tools, achieving an area under the receiver operating characteristic curve of 0.942. The model demonstrated strong generalizability across multiple external datasets and showed consistency with real-world molecular treatment proposals. MARiO offers a robust and clinically relevant approach for missense variant pathogenicity assessment by integrating variant- and patient-level features and serves as a valuable tool to support clinical decision-making.

## Introduction

Cancer is a disease driven by alterations in the genome. Therefore, targeting cancer somatic variants is an effective treatment strategy [1]. For example, patients with lung cancer who have epidermal growth factor receptor (*EGFR*) mutations have responded to EGFR tyrosine kinase inhibitors [2], demonstrating the utility of genomically matched treatments. Moreover, advances in next-generation sequencing have enabled rapid and comprehensive analysis of cancer genomes. Consequently, comprehensive genomic profiling (CGP) tests, which assess multiple cancer-related genes from patient tumors, have been approved in many countries [3–6]. CGP is typically used to enable detailed molecular profiling in patients with limited treatment options, including those with rare cancers or who have exhausted standard therapies. Upon identification of actionable alterations via CGP, molecular tumor boards (MTBs) can propose matched treatments, facilitating the targeted therapies or enrollment in clinical trials. Advancements have been made in such precision oncology strategies, and genomically matched therapies have demonstrated clinical benefit and improved patient outcomes [7–9].

CGP is increasingly used in cancer precision medicine to identify actionable variants. However, in clinical practice, variant interpretation is sometimes challenging, as not all reported missense variants have established clinical importance. CGP often reports both established variants and variants of uncertain significance (VUS) [10]. VUS are variants with unclear clinical relevance, for which currently available evidence is insufficient for definitive clinical interpretation, making it difficult to determine their relevance to treatment decision-making. Generally, such interpretations depend on annotations by testing companies and expert opinion within MTBs, often supported by knowledge bases such as ClinVar [11]. At some institutions, advanced analyses are conducted by integrating evidence from the literature, recent clinical trials, and expert opinions to refine variant interpretation. However, these processes are resource-intensive and not scalable across institutions. Thus, the development of standardized strategies for evaluating missense variants detected by CGP remains an unmet clinical need.

Recent advances in computational tools for predicting variant effects provide a valuable approach to variant interpretation [12,13]. These *in silico* methods generate scores that estimate the functional impact of mutations by leveraging features such as amino acid conservations, protein structures, and evolutionary constraints. They enable us to evaluate rare or previously unreported variants, offering an advantage over knowledge base-dependent approaches. However, these tools have several limitations, including limited coverage of certain protein regions and domain-dependent variability in predictive performance, owing to the differences in algorithms, modeling strategies, and training data. Reliance on any single *in silico* tool could reduce the robustness of variant interpretation. To address this, several ensemble methods have been developed and shown to improve predictive performance [14,15]. However, they estimate a variant’s functional impact rather than predicting its pathogenicity. In practice, clinicians synthesize information on not only the variants themselves but also clinical parameters. They also consider variant allele frequency (VAF), mutation type (e.g., somatic or germline), tumor type, and mutational backgrounds to assess pathogenicity. Existing approaches rely solely on computational predictions and fail to integrate such patient-level features. Thus, a gap remains, with the lack of a well-established framework that combines computational functional predictions with patient-specific clinical features derived from CGP testing.

To bridge this gap, we developed MARiO (Missense Alteration Risk for Oncogenicity), a computational framework that can predict the pathogenicity of cancer missense variants. MARiO integrates *in silico* variant functional predictions with patient-level genomic features, thereby bridging molecular functional evaluation and the clinical genomic context (Fig. 1). In this study, we used real-world CGP datasets to identify informative features, developed a machine learning model for pathogenicity prediction, and validated its performance across external datasets. MARiO has three key features. First, the model provides comprehensive and robust predictions by assembling multiple *in silico* tools rather than relying on a single model. Second, MARiO enables variant interpretation that reflects clinical practice by incorporating patient-specific variables (e.g., VAF). Third, although missing data frequently limit the evaluation of real-world clinical datasets, MARiO overcomes this challenge by using machine learning methods that tolerate missingness. Collectively, we present a new modeling framework with improved performance for evaluating the cancer variant pathogenicity. MARiO is designed to be accessible to a wide range of users, including non-experts, and has the potential to advance cancer precision medicine through improved interpretation of cancer missense variants.

**Figure 1.**
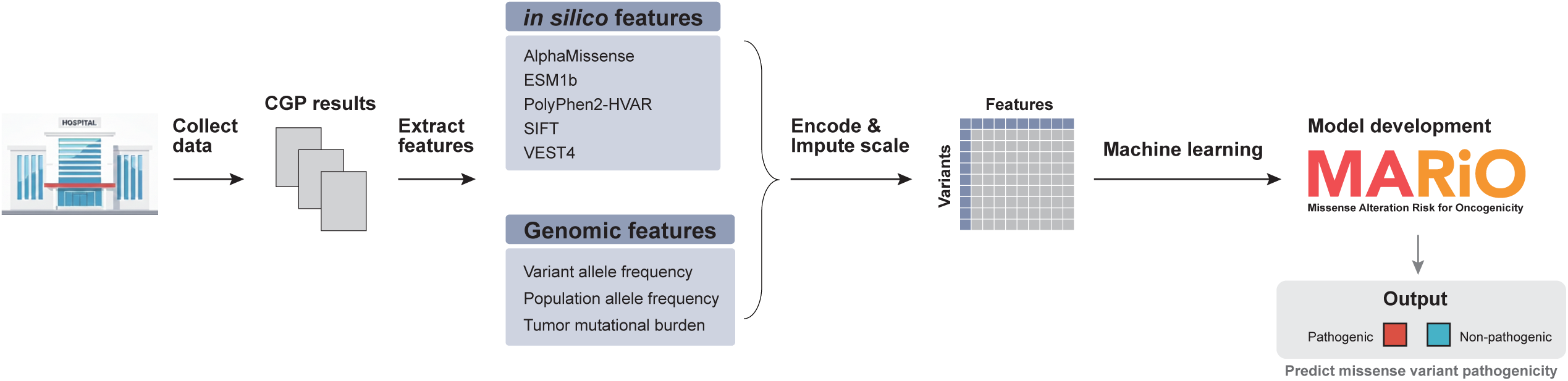
Overview of the study design. Abbreviations: CGP, comprehensive genomic profiling

## Material and methods

### Data collection

To prepare a dataset of missense variants, CGP results were retrospectively collected. We included patients whose CGP results were discussed by the MTB at the Institute of Science Tokyo (ISCT) Hospital from September 2017 to March 2024. Four CGP assay types were used in this study: FoundationOne CDx (F1) [3], FoundationOne Liquid CDx [4], OncoGuide NCC Oncopanel System [5], and Guardant360 CDx [6]. From the CGP results, missense variants were extracted for subsequent analysis. For liquid biopsy samples, variants in genes potentially arising from clonal hematopoiesis of indeterminate potential (CHIP) [16] were excluded to ensure analytical consistency (Method S1).

### Feature selection

To identify informative features for our predictive model, we evaluated candidate features based on their association with variant pathogenicity. The annotations provided by testing laboratories served as the basis for our binary labels, comprising “known pathogenic variant” and “VUS.” Following previous studies, we formulated a binary classification task by treating pathogenic variants as positive and VUS as negative [15,17] (Method S2). We then evaluated the associations between candidate features and variant pathogenicity to select features for model development. Candidate features were grouped into two categories: *in silico* variant functional prediction features and patient-level genomic features.

### *in silico* variant functional predictions

The *in silico* functional prediction features included scores from the following tools: AlphaMissense [18], ESM1b [19], EVE [20], PolyPhen2-HVAR [21], SIFT [22] and VEST4 [23]. These tools were selected because they represent either recently developed high-performance methods (AlphaMissense, ESM1b, and EVE) or widely used established tools (PolyPhen2-HVAR, SIFT, and VEST4). To assess their utility as predictive features for variant pathogenicity, we evaluated the performance of each tool in distinguishing pathogenic variants from VUS using receiver operating characteristic (ROC) and precision-recall (PR) curves.

### Patient-level genomic features

Candidate genomic features for each patient included VAF, population allele frequency (PAF), tumor mutational burden (TMB), and microsatellite instability (MSI) status. To examine whether they were statistically associated with variant pathogenicity, we performed logistic regression analysis with variant pathogenicity defined as the binary outcome, coding pathogenic variants as 1 and VUSs as 0. PAF were obtained from the Genome Aggregation Database (gnomAD) [24], specifically for East Asian populations. PAF and VAF were treated as ratio-scale variables ranging from 0 to 1. TMB was analyzed as log-transformed values, and MSI status was treated as a categorical variable with three classes: 0 for microsatellite stable, 0.5 for MSI-low, and 1 for MSI-high. For VAF, polynomial regression was used to flexibly model its association with variant pathogenicity, as VAF is influenced by multiple factors and is expected to exhibit a non-linear distribution. Given the complex nature of TMB and VAF, we further visualized their relationships with variant pathogenicity to facilitate interpretation (Method S3). Based on these analyses, the feature set was determined for inclusion in the model.

### Model development

Using the selected features, we developed models for predicting missense variant pathogenicity. However, among the features, TMB requires careful consideration because it varies across cancer types and mutated genes [25–27]; therefore, inclusion of raw TMB values may introduce systematic biases driven by these factors. To mitigate these biases and normalize TMB as model input, we introduced the “adjusted TMB,” as described below, which is a normalized TMB value according to cancer type and mutated genes.

### Cancer type- and gene-adjusted TMB modeling

To characterize dependence of the TMB on cancer type and mutated genes, we analyzed The Cancer Genome Atlas (TCGA), a large-scale database of cancer somatic mutations [28]. Missense variant data, identified via the Mutect2 pipeline, were obtained from the UCSC Xena platform [29], and variants located within CGP target genes (Method S4) within 31 TCGA cohorts (Method S5) were analyzed to determine their impact on TMB. TCGA TMB data were retrieved from the National Cancer Institute Genomic Data Commons [30,31]. We first examined the distribution of TMB across cancer types to assess cancer type effects. We also evaluated whether mutations in specific genes were associated with an increased or decreased TMB using LASSO (Least Absolute Shrinkage and Selection Operator) regression.

The analyses showed that the TMB is influenced by both cancer type and mutated genes; we therefore introduced the “adjusted TMB,” defined as the deviation of the observed TMB from its expected value given a specific cancer type and mutated gene effects. Specifically, we estimated the expected TMB for each cancer type and mutated gene and used the residual as the adjusted TMB.

To calculate the expected TMB, we modeled TMB using LASSO regression, as follows:

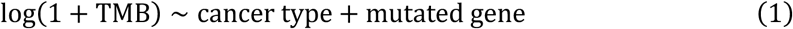

By applying an *L1* penalty to the coefficients, this model effectively selects a sparse set of genes that strongly affects the TMB while preventing overfitting.

From this model, the expected TMB for each cancer type and gene was estimated as:

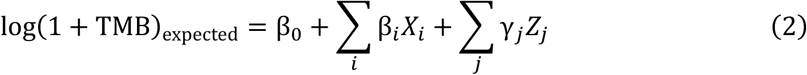

where *β_i_* and *γ_j_* represent gene-specific and cancer type-specific effects, respectively, and *X_i_* and *Z_j_* are indicator variables encoding the presence of mutation in each gene and the corresponding cancer type. For patients where cancer types do not exist in TCGA, the expected TMB was estimated using gene-specific effects only. The adjusted TMB was calculated as the difference between the observed and expected values:

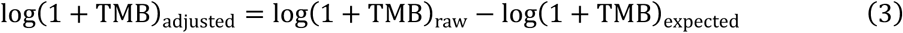

The rationale behind this calculation is to isolate the TMB variation that deviates from the baseline expected for a specific cancer type and gene mutation profile. Using this residual as an input feature, we aimed to capture the variant-specific genomic context while controlling for systematic background TMB variations. This was used as an input for all subsequent analyses.

### Machine learning model selection

Using the adjusted TMB together with all other features, we next developed predictive models of pathogenicity. To determine the optimal model construction method, we evaluated multiple machine learning models, including extreme gradient boosting (XGBoost) [32], light gradient boosting machine (LightGBM) [33], and categorical boosting (CatBoost) [34]. These methods were chosen because they can handle missing values internally, which is advantageous for real-world datasets where missing data are common.

We used nested cross-validation (CV) to evaluate model performance. Specifically, an outer 5-fold CV was used to assess performance, and an inner 3-fold CV was used for hyperparameter tuning via a randomized or grid search. We also evaluated other conventional machine learning models, including random forest, k-nearest neighbors, logistic regression, naive Bayes, and support vector machines. Because they do not allow missing values, benchmarking was performed using complete-case data. Model performance was assessed using the mean area under the ROC curve (AUROC) across outer folds. Detailed procedures are described in Method S6.

### Development and interpretation of MARiO

Based on the results, we selected the best method for final model construction. Using the best hyperparameters identified via nested CV, we trained the final model on the entire CGP dataset. The model, named Missense Alteration Risk for Oncogenicity (MARiO), provides assessment of missense variant pathogenicity. The optimal decision threshold for binary classification was determined using the Youden index, based on predictions obtained via 5-fold CV.

To interpret the model, feature contributions to the model’s predictions were evaluated using SHapley Additive exPlanations (SHAP) analysis [35]. An ablation study was performed to validate the need for the selected feature sets. Analyses were conducted using Python 3.11 with scikit-learn (v1.5.1), xgboost (v2.0.3), lightgbm (v4.4.0), catboost (v1.2.7), and shap (v0.46.0).

### Model validation using external databases

The performance of MARiO was validated using three independent external datasets: TCGA [28], Cancer Hotspots [36,37], and Memorial Sloan Kettering Cancer Center’s Precision Oncology Knowledge Base (OncoKB) [38,39].

### External validation in TCGA

MARiO, which was trained on CGP data, was evaluated on TCGA dataset. Missense variants in CGP target genes (Method S4) across 31 TCGA cohorts (Method S5) were analyzed, and a MARiO score was calculated for each variant. To benchmark MARiO’s performance, its predictions were compared against the ground-truth from ClinVar, alongside evaluations of the established *in silico* tools mentioned previously. For this validation, only variants with definitive ClinVar labels (i.e., “Pathogenic/Likely Pathogenic” as positive and “Benign/Likely Benign” as negative) were included; variants with conflicting interpretations or those lacking annotations were excluded to ensure rigorous evaluation. To assess potential cancer type-dependent differences in performance, confusion matrices and additional metrics were evaluated, stratified by cancer type.

### External validation using Cancer Hotspots

Cancer Hotspots is a curated catalog of statistically significant “hotspot mutations” in cancer. We used its classifications as ground-truth to evaluate the MARiO’s performance. Among variants registered in Cancer Hotspots, we analyzed those located in CGP-target genes (Method S5). The distribution of MARiO scores was compared between hotspot and non-hotspot variants.

### External validation in OncoKB

OncoKB, a database of functional annotations for cancer-associated mutations, offers two types of annotation: “Annotation” (variant’s oncogenicity) and “Function” (variant’s functional impact). MARiO was applied to CGP variants, and the resulting scores were evaluated for consistency with OncoKB annotations.

### Model evaluation using real-world treatment-related information

To assess the clinical utility of MARiO, we performed retrospective validation by examining the relationship between MARiO scores and real-world treatment recommendations. This information was obtained from MTB discussions of CGP results for patients who underwent F1 testing (2019–2024) at ISCT Hospital. Each variant was classified into one of four labels: (1) an approved drug was recommended and administered, (2) an investigational drug (e.g., clinical trials, off-label use, or unapproved drugs in Japan) was recommended and administered, (3) a drug was recommended but was not administered, or (4) no therapeutic recommendation was provided. Exclusion criteria were lack of consent to participate in the research, death before results disclosure, and loss to follow-up. Treatment-related data were analyzed for associations with MARiO scores.

## Results

### Patient and variant characteristics

From September 2017 to March 2024, 1274 patient cases were discussed by the MTB at ISCT Hospital. Among them, 1271 patients harbored at least one missense variant. Among them, 1021 patients (80.3%) were tested using F1, 166 patients (13.1%) using FoundationOne Liquid CDx, 62 patients (4.9%) using NCC Oncopanel System, and 22 patients (1.7%) using Guardant360 CDx (Table 1). In total, 10,687 missense variants were collected (Tables S1, S2). Of these, 462 CHIP-suspected variants were excluded, leaving 10,225 variants for subsequent analyses (Fig. S1).

**Table 1.**
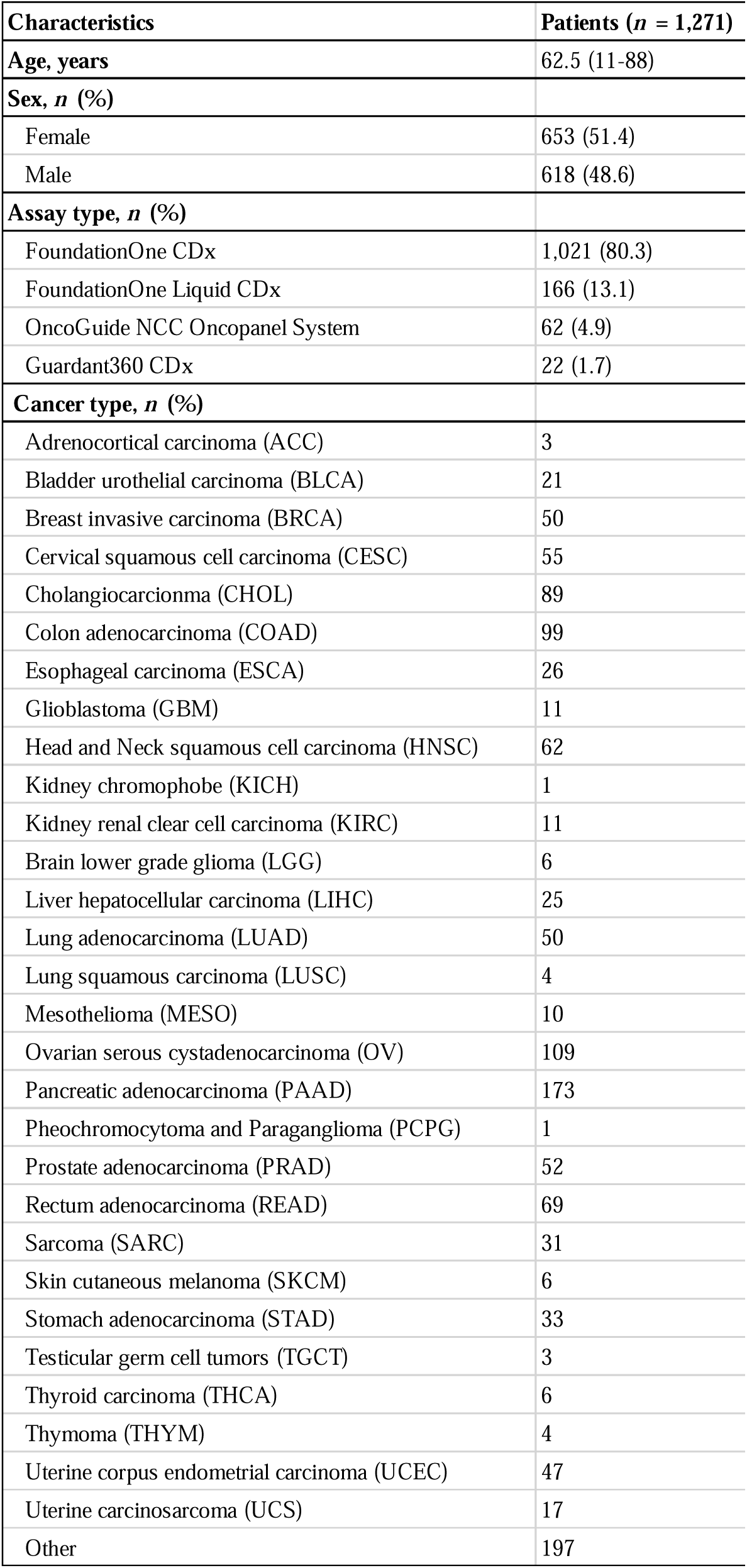
Patient characteristics.

### Feature characterization and selection

We examined whether *in silico* functional prediction scores could predict pathogenic missense variants. All tools demonstrated the ability to distinguish pathogenic variants from VUS, with AlphaMissense performing the best; this suggested their utility for pathogenicity assessment (Fig. 2a). We then assessed the proportion of missing values for each tool (Fig. 2b, Table S3). Notably, EVE failed to generate scores for approximately half the variants, resulting in a substantially higher rate of missing values compared with the other tools. Based on these observations, we selected five *in silico* prediction tools, excluding EVE, as features to include.

**Figure 2.**
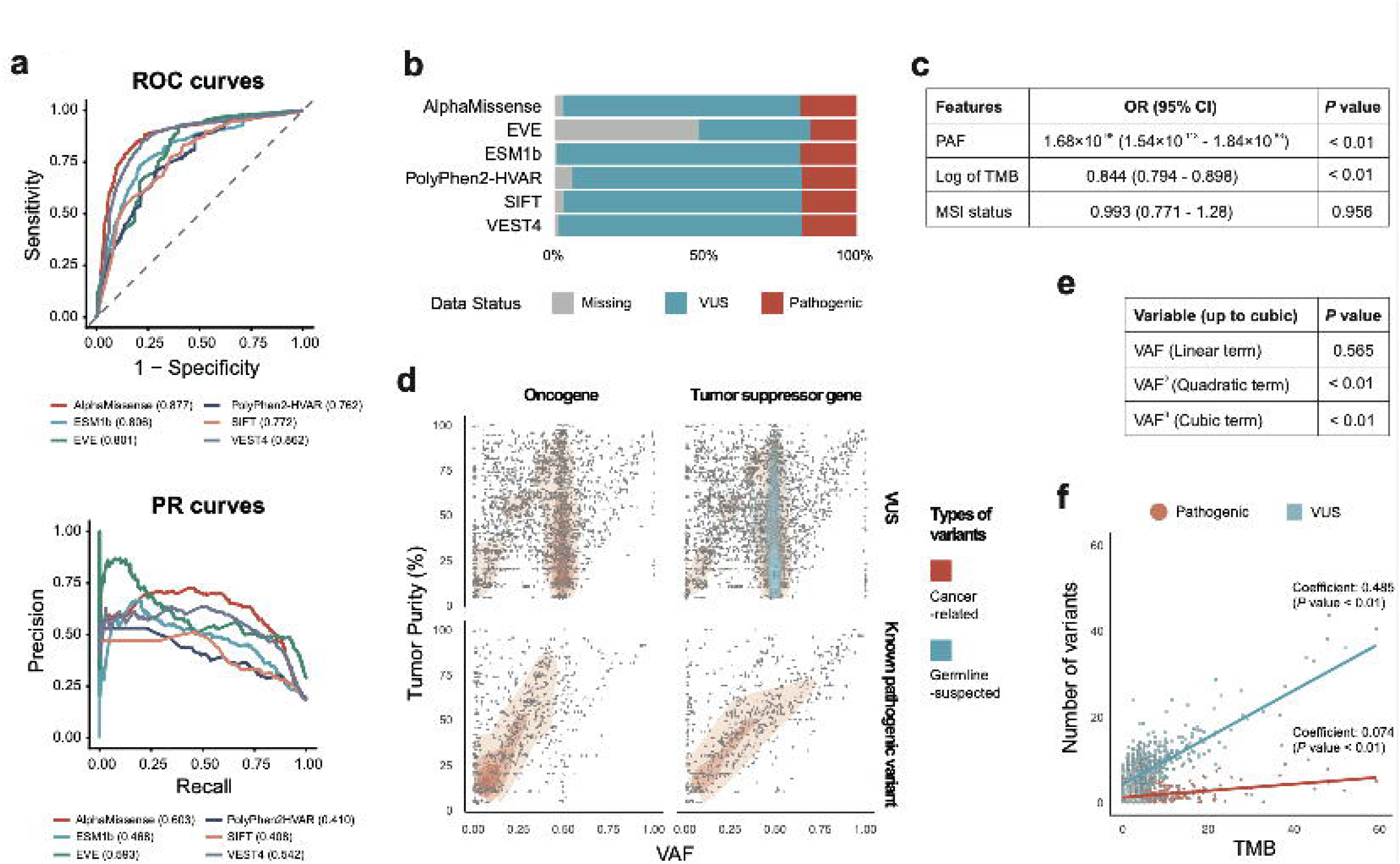
Evaluation of *in silico* tools and characterization of patient genomic features. **a** ROC and PR curves for AlphaMissense, EVE, ESM1b, PolyPhen2-HVAR, SIFT, and VEST4. Values in parentheses indicate the AUROC and PR-AUC, respectively. **b** Proportion of missing predictions and annotation labels for each *in silico* tool. **c** Logistic regression analysis of patient-specific genomic features, including PAF, log-transformed TMB, and MSI status. **d** Relationship between VAF and tumor purity. Each point represents an individual variant. Panels are stratified by variant pathogenicity and gene function. The color map indicates variant categories: red, variants in cancer-related genes (oncogenes or tumor suppressor genes); blue, germline-suspected variants defined as variants with population PAF ≥ 1% in gnomAD. **e** Polynomial regression analysis modeling the relation between VAF and variant pathogenicity. **f** Relationship between TMB and variant burden by variant category. Scatter plots show the number of pathogenic variants and VUS per patient (y-axis) against patient-level TMB (x-axis). Blue indicates VUS and red indicates pathogenic variants.

Next, we evaluated the associations between genomic features (PAF, TMB, and MSI status) and variant pathogenicity. No significant association was observed for MSI status whereas PAF and TMB were significantly associated with variant pathogenicity (Fig. 2c). Variant pathogenicity showed a strong inverse association with PAF (OR 1.68 × 10^−98^, 95% 1.54 × 10^−113^–1.84 × 10^−83^, *P* < 0.01) and an inverse association with TMB (OR 0.844, 95% CI 0.794–0.898, *P* < 0.01).

Among these features, the PAF’s importance can be readily explained from a biological perspective. Pathogenic variants tend to have low PAFs, which is because deleterious single nucleotide polymorphisms (SNPs) are removed from the population through purifying selection.

VAF exhibited a complex distribution, which differed according to variant type (pathogenic or VUS) and gene class (tumor suppressor gene or oncogene) (Fig. 2d). For VUS, variants were frequently observed at a VAF of approximately 50%, consistent with a germline-like distribution. By contrast, pathogenic variants clustered around VAF = purity and VAF = 1/2 purity. Pathogenic variants in oncogenes were accumulated around VAF = 1/2 purity, consistent with heterozygous activating mutations. However, tumor suppressor gene variants were accumulated at both VAF = purity and VAF = 1/2 purity. In this case, VAF = purity variant likely reflects a loss of heterozygosity event, and VAF = 1/2 purity would represent compound heterozygosity [40] (the mutations in different codons of homologous genes) or multiple heterozygous aberrations in the same signaling pathway (like BRCAness) [41]. These observations suggest that VAF harbors biologically meaningful information, but the relationship is not linear. To model this complex association, we assessed VAF using polynomial regression, which enables non-linear modeling. This revealed that the second- and third-degree VAF terms were significantly associated with pathogenicity (Fig. 2e), suggesting a non-linear contribution of VAF to variant pathogenicity. Collectively, these results indicate that VAF is an informative feature for prediction, although its effect is non-linear and context dependent.

While TMB was associated with pathogenicity in our rs, the biological basis has not been well characterized. We therefore examined the association between TMB and variant counts per individual. Increasing TMB was associated with a larger increase in the number of VUS than that in pathogenic variants (Fig. 2f). Two-way analysis of variance revealed a significant interaction effect between TMB and variant counts by type (*P* < 0.001; Table S4), suggesting that increases in TMB were disproportionately driven by VUS. This means that in high-TMB tumors, the relative pathogenic contribution of individual variants is reduced, consistent with the observed negative association between TMB and variant pathogenicity.

Based on these findings, eight variables were selected for model development: five in silico variant functional predictions, VAF, PAF, and TMB.

### Model development

Using the selected features, we developed a predictive model for missense variant pathogenicity. Among the candidate features, TMB is known to be influenced by multiple biological factors, which could introduce biases if raw values are directly used. Therefore, prior to model construction, we characterized TMB and performed TMB adjustment.

### Cancer type- and gene-adjusted TMB modeling

To characterize the association between TMB and cancer type, we visualized the distribution of TMB across cancer types using TCGA data. TMB showed substantial heterogeneity across cancer types (Fig. 3a). The association was evaluated using LASSO regression, and several genes exhibited positive or negative effects on the TMB (Fig. 3b, Table S5), suggesting that mutations in specific genes are associated with shifts in the TMB.

**Figure 3.**
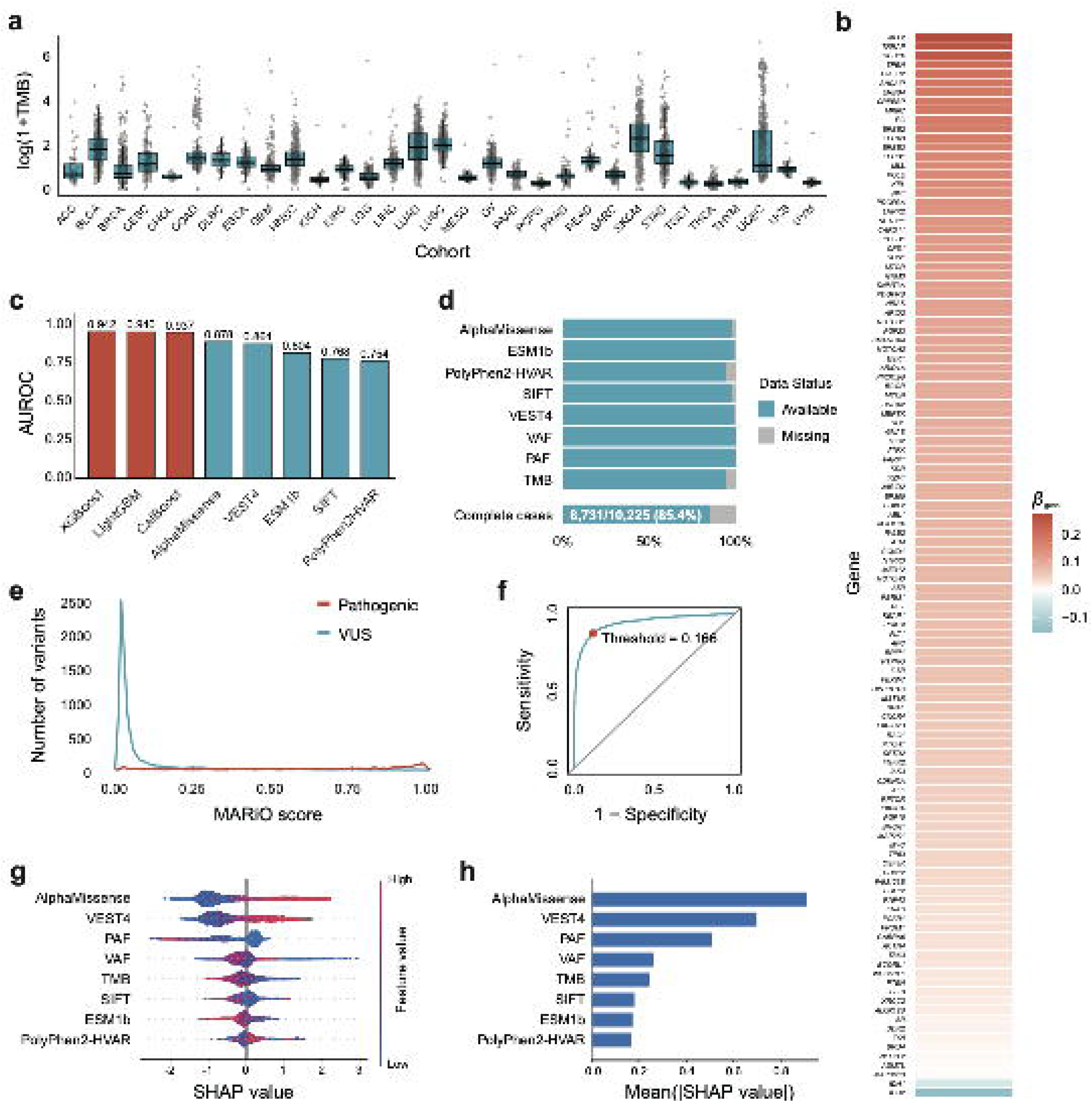
TMB modeling for feature adjustment and development of the MARiO model. **a** Distribution of TMB across cancer types (see abbreviations in Table 1) in TCGA. **b** Gene-specific coefficients from regression modeling, with TMB as the response variable. The heatmap shows genes with non-zero coefficients. **c** Comparison of predictive performance between machine learning models (XGBoost, LightGBM, and CatBoost; red) and existing *in silico* tools (blue), assessed using the AUROC. **d** Proportion of missing values for each feature. Complete-case variants accounted for 8731 of 10,225 variants. **e** Distribution of MARiO scores across CGP variants. **f** ROC curve of the MARiO model. The optimal decision threshold (0.166) was determined using the Youden index. **g** Beeswarm plot of SHAP values. The x-axis represents the contribution to the model output. **h** Bar plot of mean absolute SHAP values. Higher values indicate stronger feature contributions to the model predictions.

Considering these effects, we calculated the expected TMB for each cancer type and mutated gene combination (Table S6). For cancer types not represented in TCGA, gene-specific expected TMB values were determined. These expected TMB values were used to calculate the adjusted TMB, which served as an input feature for downstream analysis.

### Comparison of machine learning models

Using the adjusted TMB and other features, we compared the three machine-learning models. XGBoost performed the best, with a mean AUROC of 0.942 (Fig. 3c). All models outperformed conventional tools. These models, which can inherently handle missing data, have an advantage over other machine-learning methods that do not tolerate missingness. This is because real-world datasets contain a substantial proportion of missing values (Fig. 3d), so models that can handle missing data would fail to evaluate approximately one in every several patients. We also compared these models with other conventional models that cannot handle missing data, on a complete case (8731 variants). In this setting, gradient boosting methods (XGBoost, LightGBM, and CatBoost) consistently ranked as the top-performers, followed by other methods such as random forest (Fig. S2). An ablation study revealed that the base model showed the highest AUROC among the tested feature sets (Fig. S3).

### Development and interpretation of MARiO

Based on these results, we developed MARiO, a machine learning model for predicting missense variant pathogenicity in patients with cancer. MARiO integrates *in silico* variant functional evaluations and patient-specific genomic features via XGBoost, enabling predictions by leveraging both annotation independent functional predictions and individual mutational contexts. The MARiO scores effectively distinguished pathogenic variants from VUS (Fig. 3e). We also assessed this using SHAP (Fig. 3f, 3g) to interpret the contribution of each feature, and revealed that AlphaMissense and VEST4 contributed strongly and positively to prediction whereas PAF had a negative impact on the model output. The pathogenicity classification threshold was set at 0.166, determined by maximizing the Youden index on the ROC curve (Fig. 3h); this was applied to all subsequent performance evaluations and comparative analyses.

### Model validation using external databases

#### Validation on ClinVar labels using TCGA data

To assess generalizability of the model, MARiO was benchmarked against conventional computational tools using TCGA variants. MARiO outperformed conventional tools with higher accuracy and recall (Fig. 4a, 4b) and maintained consistent performance across different cancer types (Fig. S4). Additionally, MARiO produced fewer false negatives than other tools, leading to a reduced risk of overlooking truly pathogenic variants, which is important in clinical settings. These results support MARiO’s generalizability to independent external datasets and demonstrate its robustness across heterogeneous cancer types. As a supplementary analysis, we compared MARiO with a model excluding the adjusted TMB; MARiO outperformed this ablated model, demonstrating the efficacy of the adjusted TMB as a feature (Fig. S5).

**Figure 4.**
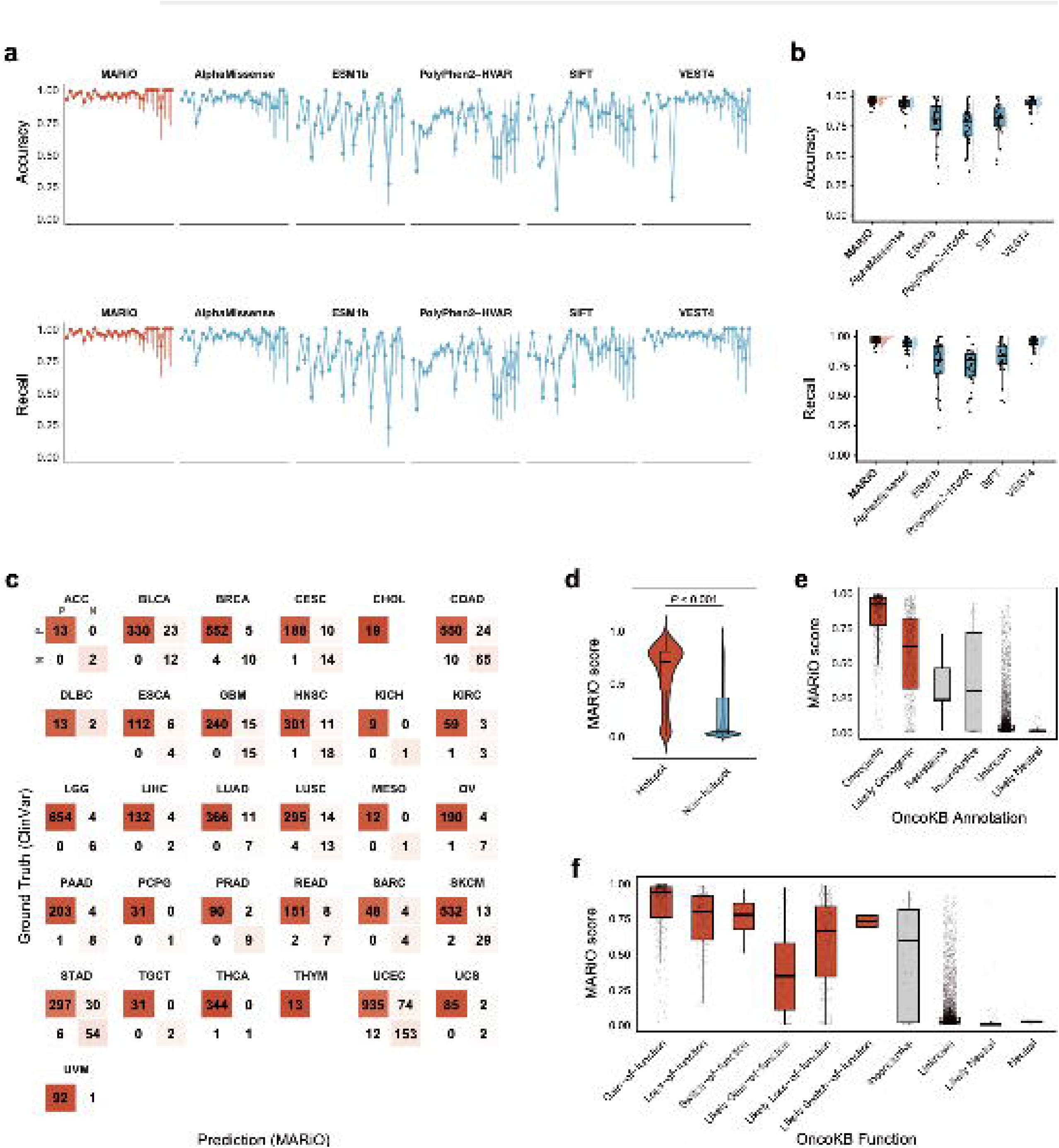
Validation of MARiO using external datasets. **a** Line plots showing the accuracy and recall of MARiO and *in silico* tools across cancer types. Each point represents the performance metric for each cancer type, with vertical bars indicating 95% confidence intervals calculated using the Wilson’s score method. Cancer types are ordered from left to right by decreasing sample size. The ordering of cohorts is as follows: UCEC, LGG, COAD, SKCM, BRCA, STAD, LUAD, BLCA, THCA, HNSC, LUSC, GBM, PAAD, CESC, OV, READ, LIHC, ESCA, PRAD, UVM, UCS, KIRC, SARC, TGCT, PCPG, CHOL, ACC, DLBC, MESO, THYM, KICH. For cancer type abbreviations, see Table 1. **b** Raincloud plots showing cancer-type-specific accuracy or recall for MARiO and *in silico* tools. Each point represents a cancer type, with density and boxplots summarizing the distribution across cancer types. MARiO’s accuracy and recall were significantly higher than those of conventional tools (Wilcoxon rank-sum test, Benjamini–Hochberg adjusted *P* < 0.05). **c** Confusion matrix for MARiO, evaluated on TCGA missense variants with ClinVar annotations. Variants were classified as pathogenic (“Pathogenic” and “Likely Pathogenic”) or benign (“Benign” and “Likely Benign”), and the labels were used as the ground truth. **d** Distributions of MARiO scores for hotspot and non-hotspot variants based on Cancer Hotspots. MARiO scores were significantly higher for hotspot variants than for non-hotspot variants (Mann–Whitney U test, *P* < 0.001). **e** Distribution of MARiO scores stratified by OncoKB oncogenicity annotations, which classify variants according to their cancer-driving potential. **f** Distribution of MARiO scores stratified by OncoKB functional impact annotations, which categorize variants by curated functional effects.

#### Validation on hotspot labels in Cancer Hotspots

Cancer Hotspots harbored 3166 missense variants as hotspots, of which 2806 were variants in genes analyzed in CGP. MARiO scores were significantly higher for hotspot variants compared with non-hotspot variants (*P* < 0.001; Fig. 4c). Gene-level analyses revealed that this trend was consistent across most genes, although some exceptions were observed (Fig. S6). For instance, in *APC*, no significant difference in MARiO scores was found between hotspot and non-hotspot missense variants. Overall, MARiO scores were consistently elevated for hotspot variants across most genes, demonstrating the consistency of MARiO in identifying oncogenic hotspot mutations.

#### Validation on OncoKB annotations

Variants annotated as oncogenic in OncoKB exhibited higher MARiO scores whereas most neutral- or unknown-annotated variants were skewed toward lower scores (Fig. 4d). Similarly, variants labeled as functionally changing (“Gain-of-function,” “Likely Gain-of-function,” “Loss-of-function,” “Likely Loss-of-function,” “Switch-of-function,” and “Likely Switch-of-function”) showed higher MARiO scores than variants annotated as neutral or unknown (Fig. 4e). These results showed that MARiO scores are consistent with curated functional and oncogenic annotations in OncoKB, supporting the biological relevance of its predictions.

### Model evaluation using real-world treatment-related information

Finally, we assessed MARiO’s clinical utility using real-world patient outcome data from our hospital. At ISCT Hospital, 420 patients underwent F1 testing from October 2019 to March 2024 (Fig. S7). Six individuals were excluded: three did not provide informed consent, two died before results disclosure, and one returned to their home country, making follow-up impossible. Across the 414 patients, 3319 missense variants were identified, of which 289 variants were recommended for some treatment options. Variants with candidate drug recommendations consistently exhibited a skewed distribution toward higher MARiO scores; no such skewness was observed for variants without candidate drugs (Fig. 5a, 5b). This suggests that variants with treatment proposals have higher scores, demonstrating the model’s utility in predicting variants with actionable therapeutic agents.

**Figure 5.**
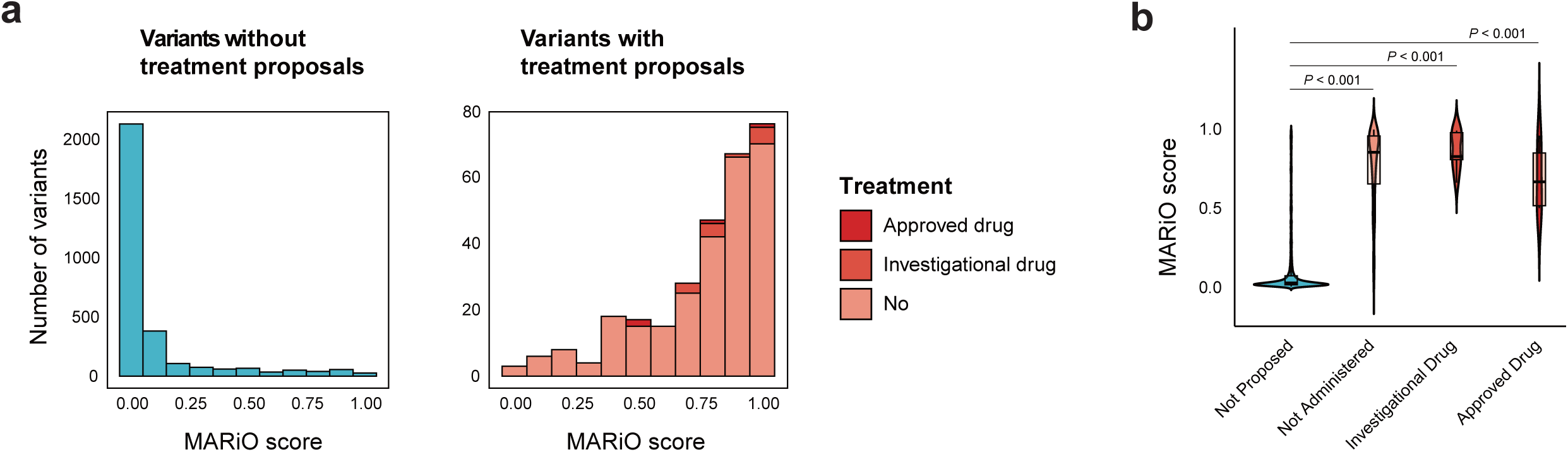
Validation of MARiO using treatment-related information. **a** Distribution of MARiO scores for missense variants stratified by the presence or absence of treatment recommendations. The left panel shows variants without treatment options (n = 3030), and the right panel shows variants with at least one proposed treatment option (n = 289). Treatment categories are indicated by color: red, approved drugs proposed and administered; orange, investigational or off-label drugs recommended and administered; pale orange, candidate drugs identified but not administered. **b** Violin plots showing the distribution of MARiO scores for variants stratified by treatment recommendations Treatment categories are indicated by color (red, approved drugs; orange, investigational or off-label drugs; pale orange, candidate drugs). Pairwise comparisons between no-treatment group and each treatment category were performed by the Mann–Whitney U test.

## Discussion

In this study, we first analyzed real-world CGP data and explored features to predict pathogenicity of missense variants and identified *in silico* functional predictions and genomic features related to variant pathogenicity. Based on these findings, we developed a new variant pathogenicity prediction model, MARiO, by integrating *in silico* functional predictions and genomic features with machine learning. This model significantly outperformed conventional tools, showed generalizability to external datasets, and demonstrating its clinical utility in treatment proposals.

MARiO integrates several features that capture distinct biological perspectives. First, the contribution of *in silico* functional predictions suggests the effectiveness of computational approaches in assessing the variant-level functional impact. Second, the negative contribution of PAF was observed, which could be explained by purifying selection. Third, VAF could provide complementary information from genomic perspectives. As described in the results, VAF was differentially distributed depending on whether variants were germline or somatic, oncogenic or tumor-suppressive, and pathogenic or non-pathogenic. These observations indicate that VAF encodes biologically meaningful information about the role of variants in cancer. Furthermore, VAF may reflect clonal dominance within tumors. Driver mutations are typically subject to positive selection during oncogenesis and therefore tend to become clonal events with higher VAF. Consequently, VAF reflects the functional relevance of a variant within the tumor’s evolutionary context. Finally, our results revealed the contribution of TMB. In TMB-high cases, numerous passenger mutations reduce the relative contribution of any single variant to tumor. By contrast, TMB-low tumors are often driven by a limited number of strong driver mutations, so individual variants may exert a greater biological impact. Interestingly, the presence of mutations in *IDH1/2* was associated with lower TMB; this trend has been reported in previous studies [26,42,43]. Although further study is needed to explore the reasons, one possible explanation is that these potent driver mutations can promote oncogenesis with minimal additional mutational burden. By contrast, several patterns were observed among genes associated with TMB-high tumors. One group includes genes known to drive hypermutation, such as *POLE* and *POLD1*. Another pattern is characterized by an accumulation of variants in genes frequently classified as VUS, such as *SPEN*, *TRRAP*, and *KMT2D*. Notably, these are all exceptionally large genes, which are prone to accumulating random passenger mutations with increased TMB, thereby inflating the total variant count without necessarily increasing pathogenic drive. This inverse relationship between TMB and individual variant significance could explain the contribution of TMB in our model.

In model development, we introduced the adjusted TMB to account for cancer type- and gene-specific effects on TMB. For example, cancers such as melanoma and non-small cell lung cancer are known to exhibit a globally elevated TMB at the cohort level, and mutations in DNA replication and repair-associated genes, like *POLE*, are well established as dramatically increasing the TMB [44]. By adjusting for these background effects, the model can use the mutational context in a biologically meaningful way, with TMB providing information beyond cancer type- and gene-specific background effects.

In validation analysis using Cancer Hotspots, MARiO scores were generally higher for hotspot variants, with *APC* being an exception. This likely reflects *APC*’s unique molecular patterns, where nonsense variants are more prevalent than missense variants [45,46]. Such gene-specific distributions of variant types intrinsically influence the model’s predictive performance.

This study has several limitations. The first is data bias because all data were from patients with cancer, which potentially introduces selection bias. Additionally, Japanese insurance coverage for CGP is restricted to patients with no remaining standard therapies or rare cancers, so most variants are derived from late-stage cancers. This could introduce molecular patterns that may differ from those in early-stage cancers, suggesting that the current model may not fit cancers in the early stage. Additionally, potential biases may exist within the datasets used for training. We used gnomAD SNP data as the PAF reference, but SNP patterns vary across populations. Model retraining with appropriate data could help address these issues.

Another limitation lies in the ground-truth labels used in this study. We relied on the company’s annotations and used them as the ground truth, but they may not always be definitive. For instance, variants classified as VUS could be pathogenic but remain unconfirmed owing to a lack of experimental validation. Therefore, using more appropriate ground truth could contribute to improving the model.

Finally, this study was limited to retrospective validation. Prospective validation is desirable to assess the clinical impact of our framework. For example, MARiO may predict some variants classified as VUS in existing databases as being pathogenic, and by confirming that such variants are truly oncogenic, we would further support our method. Indeed, we experienced a case in which only VUS and no established pathogenic variants were reported in CGP, but one of these VUS was predicted to be pathogenic by MARiO. In this case, a matched drug was administered for the predicted pathogenic variant, and tumor shrinkage was observed. We are currently accumulating such cases, and these findings will be reported in future case reports.

In summary, we developed MARiO, a new predictive model for missense variant pathogenicity using real-world CGP data. MARiO integrates multiple *in silico* predictions with patient-level genomic features, enabling holistic assessment that outperformed existing tools. Our study provides a framework for improving the interpretation of cancer missense variants and advancing precision cancer medicine.

## Acknowledgments

We thank Junko Yokobori and Mika Ohki for their excellent coordination and management of patient information as cancer genome medical coordinators, and Akiko Tobe and Takamasa Fujita for the interpretation and annotation of variant data. Our sincere thanks go to Yumi Kobayashi for her contribution in database curation, and to Rieko Furuya, Akiko Noguchi, Tomoko Murata, Ryoko Seki for their administrative support and patient care. We also thank Analisa Avila, MPH, ELS, of Edanz for editing a draft of this manuscript. Finally, we thank all the patients for their cooperation with this study.

## Disclosure

### Ethics Statement

- Approval of the research protocol by an Institutional Reviewer Board: The study was approved by the institutional review boards of ISCT (M2024-62). All studies were performed in accordance with the relevant guidelines and regulations.
- Informed Consent: Passive consent was obtained from all patients included in the study through an opt-out consent procedure.
- Registry and the Registration No. of the study/trial: N/A
- Animal Studies: N/A

### Author contributions

HN, TK, and SI designed the study and conducted data curation, analysis, writing the original manuscript draft, and manuscript review and editing. NI, SA, MM, and FM contributed to data curation, interpretation, and manuscript review and editing. HN and TK were responsible for statistical analysis. All authors read and approved the final version of the manuscript.

## List of Supporting Information

Additional file 1: Supplementary_Materials.pdf

Includes all supplementary methods (Method S1 to S6), all supplementary figures (Fig S1 to S7), and supplementary tables (Table S1 to S4).

Additional file 2: TableS5.xlsx

Includes one supplementary table (Table S5).

## Abbreviations

AUROC: area under the receiver operating characteristic curve
CGP: comprehensive genomic profiling
MARiO: Missense Alteration Risk for Oncogenicity
MSI: microsatellite instability
OncoKB: Memorial Sloan Kettering Cancer Center’s Precision Oncology Knowledge Base
PAF: population allele frequency
PR: precision recall
PR-AUC: area under the PR curve
ROC: receiver operating characteristic
TCGA: The Cancer Genome Atlas
TMB: tumor mutational burden
VAF: variant allele frequency
VUS: variant of uncertain significance

