## Supplementary Materials for "MARiO: predicting cancer variant pathogenicity by integrating *in silico* evaluation and patient-level mutational contexts"

**Method S1.** Removal of CHIP-Suspected Variants for Liquid Biopsy Samples
**Method S2.** Grouping of Annotation Labels for Variant Pathogenicity
**Method S3.** Characterization of Genomic Features for Feature Selection
**Method S4.** List of Target Genes Analyzed in Comprehensive Genomic Profiling (CGP) Tests
**Method S5.** TCGA Cohorts Included in This Study
**Method S6.** Machine Learning Model Development and Evaluation

**Table S1.** Patient Counts by Institution
**Table S2.** Variant Counts by Annotation Category and Test Type
**Table S3.** Annotation Categories and Missing Scores of *In Silico* Tool
**Table S4.** Two-way ANOVA Results for TMB and Variant Type

**Figure S1.** Variant Selection Workflow
**Figure S2.** Model Performance on Complete-Case Data
**Figure S3.** Ablation Study of the Machine Learning Model
**Figure S4.** Performance of *In Silico* Tools on TCGA Variants Based on ClinVar Labels
**Figure S5.** Comparison of MARiO and a Model Without Adjusted TMB using TCGA Variants Based on ClinVar Labels
**Figure S6.** Validation of MARiO Scores Using the Cancer Hotspots Catalog
**Figure S7.** Cohort for Treatment-Related Analyses

**Supplementary References**

#### Method S1. Removal of CHIP-Suspected Variants for Liquid Biopsy Samples

To ensure analytical consistency, variants potentially arising from clonal hematopoiesis of indeterminate potential (CHIP) were excluded from the analysis. CHIP-associated variants can be detected in circulating cell-free DNA and may confound the interpretation of tumor-derived variants. Therefore, we removed variants in genes with a high likelihood of being associated with CHIP, based on previous reports [1].

Among the liquid biopsy results, variants in the following genes were excluded from the analysis owing to a high likelihood of being associated with CHIP: *AR, ARID2, ASXL1, ATM, CHEK2, DNMT3A, IDH2, IRS1, JAK2, MLL2, MPL, NF1, GNAS, ROS1, TET2*, and *TP53*.

#### Method S2. Grouping of Annotation Labels for Variant Pathogenicity

Annotation labels provided by testing laboratories were harmonized according to predefined subgrouping rules. For each assay, the following subgrouping criteria were applied.

**1. FoundationOne CDx (F1) and FoundationOne Liquid CDx (F1L)**

Variants labeled as "known" or "likely" in F1 and F1L were grouped as known pathogenic variants. Variants annotated as "VUS" (variants of unknown significance) were classified as VUS.

**2. OncoGuide NCC Oncopanel System (NOP)**

Variants labeled as "actionable" were categorized as known pathogenic variants, and those annotated as "VUS" were classified as VUS. By contrast, variants annotated as "incidental" were excluded from the analysis, as this designation applies to genes rather than individual variants. Such genes are associated with high disclosure recommendations owing to their medical actionability within the context of germline mutations.

**3. Guardant360 CDx (G360)**

Variants labeled as "level 1" and "level 2" were classified as known pathogenic variants, with level 1 encompassing *KRAS* G12C (the only missense variant in this category) and level 2 covering all other known pathogenic missense variants. Variants annotated as "level 4" were grouped as VUS. Variants labeled as "level 3," corresponding to non-missense variants (e.g., fusions, copy number alterations), were excluded from the analysis.

#### Method S3. Characterization of Genomic Features for Feature Selection

##### 1. Analysis of VAF

Variant allele frequency (VAF) was analyzed separately owing to its anticipated non-linear distribution. To characterize its behavior, we first visualized the distribution of VAF in relation to tumor purity and functional gene annotations.

Functional gene annotations and the putative germline status were overlaid on the distribution plots of VAF. These annotation labels were obtained as follows.

a) Functional gene annotations: oncogenes and tumor suppressor genes

Annotations for oncogenes and tumor suppressor genes were obtained from OncoKB, a database providing functional annotations for cancer-associated mutations [2.3].

b) Germline-suspected variants

Germline-suspected variants were defined based on population allele frequency data from the Genome Aggregation Database, with variants exhibiting a population allele frequency of ≥ 1% classified as potential germline variants.

In addition to visualization, polynomial regression analysis including terms up to the third-degree was performed to model the nonlinear relationship between VAF and variant pathogenicity.

##### 2. Analysis of TMB

To assess the relationship between the tumor mutational burden (TMB) and variant categories, two analyses were conducted.

First, scatter plots were generated to examine the relationship between TMB and the number of variants per patient. For each patient, TMB was plotted on the x-axis, and the number of variants of a given category (VUS or pathogenic variants) was plotted on the y-axis, resulting in two data points per patient. In other words, scatter plots were generated for each variant category to examine whether the relationship between TMB and number of variants differed between VUS and pathogenic variants.

Second, we fitted a linear model including an interaction term between the TMB and variant category to evaluate whether the relationship between TMB and variant count differed by variant category. Variant category (VUS or pathogenic variants) was treated as a binary factor, and the number of variants per patient was used as the response variable. The interaction effect between TMB and variant category was assessed using analysis of variance (ANOVA).

**Method S4. List of Target Genes Analyzed in Comprehensive Genomic Profiling (CGP) Tests**

The genes targeted for analysis in F1, F1L, NOP, and G360 were analyzed in the validation study. The following genes were included:

*ABL1, ACVR1B, AKT1, AKT2, AKT3, ALK, ALOX12B, AMER1, APC, AR, ARAF, ARFRP1, ARID1A, ASXL1, ATM, ATR, ATRX, AURKA, AURKB, AXIN1, AXL, BAP1, BARD1, BCL2, BCL2L1, BCL2L2, BCL6, BCOR, BCORL1, BRAF, BRCA1, BRCA2, BRD4, BRIP1, BTG1, BTG2, BTK, CALR, CARD11, CASP8, CBFB, CBL, CCND1, CCND2, CCND3, CCNE1, CD22, CD274, CD70, CD79A, CD79B, CDC73, CDH1, CDK12, CDK4, CDK6, CDK8, CDKN1A, CDKN1B, CDKN2A, CDKN2B, CDKN2C, CEBPA, CHEK1, CHEK2, CIC, CREBBP, CRKL, CSF1R, CSF3R, CTCF, CTNNA1, CTNNB1, CUL3, CUL4A, CXCR4, CYP17A1, DAXX, DDR1, DDR2, DIS3, DNMT3A, DOT1L, EED, EGFR, EMSY, EP300, EPHA3, EPHB1, EPHB4, ERBB2, ERBB3, ERBB4, ERCC4, ERG, ERRFI1, ESR1, EZH2, FANCA, FANCC, FANCG, FANCL, FAS, FBXW7, FGF10, FGF12, FGF14, FGF19, FGF23, FGF3, FGF4, FGF6, FGFR1, FGFR2, FGFR3, FGFR4, FH, FLCN, FLT1, FLT3, FOXL2, FUBP1, GABRA6, GATA3, GATA4, GATA6, GID4, GNA11, GNA13, GNAQ, GNAS, GRM3, GSK3B, H3-3A, HDAC1, HGF, HNF1A, HRAS, HSD3B1, ID3, IDH1, IDH2, IGF1R, IKBKE, IKZF1, INPP4B, IRF2, IRF4, IRS2, JAK1, JAK2, JAK3, JUN, KDM5A, KDM5C, KDM6A, KDR, KEAP1, KEL, KIT, KLHL6, KMT2A, KMT2D, KRAS, LTK, LYN, MAF, MAP2K1, MAP2K2, MAP2K4, MAP3K1, MAP3K13, MAPK1, MAPK3, MCL1, MDM2, MDM4, MED12, MEF2B, MEN1, MERTK, MET, MITF, MKNK1, MLH1, MPL, MRE11, MSH2, MSH3, MSH6, MST1R, MTAP, MTOR, MUTYH, MYC, MYCL, MYCN, MYD88, NBN, NF1, NF2, NFE2L2, NFKBIA, NKX2-1, NOTCH1, NOTCH2, NOTCH3, NPM1, NRAS, NSD2, NSD3, NT5C2, NTRK1, NTRK2, NTRK3, P2RY8, PALB2, PARP1, PARP2, PARP3, PAX5, PBRM1, PDCD1, PDCD1LG2, PDGFRA, PDGFRB, PDK1, PIK3C2B, PIK3C2G, PIK3CA, PIK3CB, PIK3R1, PIM1, PMS2, POLD1, POLE, PPARG, PPP2R1A, PPP2R2A, PRDM1, PRKAR1A, PRKCI, PRKN, PTCH1, PTEN, PTPN11, PTPRO, QKI, RAC1, RAD21, RAD51, RAD51B, RAD51C, RAD51D, RAD52, RAD54L, RAF1, RARA, RB1, RBM10, REL, RET, RHEB, RHOA, RICTOR, RIT1, RNF43, ROS1, RPTOR, SDHA, SDHB, SDHC, SDHD, SETD2, SF3B1, SGK1, SMAD2, SMAD4, SMARCA4, SMARCB1, SMO, SNCAIP, SOCS1, SOX2, SOX9, SPEN, SPOP, SRC, STAG2, STAT3, STK11, SUFU, SYK, TBX3, TEK, TENT5C, TERT, TET2, TGFBR2, TIPARP, TNFAIP3, TNFRSF14, TP53, TSC1, TSC2, TYRO3, U2AF1, VEGFA, VHL, WT1, XPO1, XRCC2, ZNF217,* and *ZNF703.*

#### Methods S5. TCGA Cohorts Included in This Study

This analysis included 31 cohorts of The Cancer Genome Atlas (TCGA), as follows: ACC, adrenocortical carcinoma; BLCA, bladder urothelial carcinoma; BRCA, breast invasive carcinoma; CESC, cervical squamous cell carcinoma and endocervical adenocarcinoma; CHOL, cholangiocarcinoma; COAD, colon adenocarcinoma; DLBC, lymphoid neoplasm diffuse large B-cell lymphoma; ESCA, esophageal carcinoma; GBM, glioblastoma; HNSC, head and neck squamous cell carcinoma; KICH, kidney chromophobe; KIRC, kidney renal clear cell carcinoma; LGG, brain lower grade glioma; LIHC, liver hepatocellular carcinoma; LUAD, lung adenocarcinoma; LUSC, lung squamous cell carcinoma; MESO, mesothelioma; OV, ovarian serous cystadenocarcinoma; PAAD, pancreatic adenocarcinoma; PCPG, pheochromocytoma and paraganglioma; PRAD, prostate adenocarcinoma; READ, rectum adenocarcinoma; SARC, sarcoma; SKCM, skin cutaneous melanoma; STAD, stomach adenocarcinoma; TGCT, testicular germ cell tumors; THCA, thyroid carcinoma; THYM, thymoma; UCEC, uterine corpus endometrial carcinoma; UCS, uterine carcinosarcoma; UVM, uveal melanoma.

#### Method S6. Machine Learning Model Development and Evaluation

To select the best predictive model, multiple machine learning methods were systematically evaluated. Model performance was assessed using nested cross-validation, consisting of an outer 5-fold cross-validation loop and an inner 3-fold loop for hyperparameter optimization. Hyperparameters were tuned using either grid search or random search, depending on the model. The area under the receiver operating characteristic curve (AUROC) was used as the optimization metric during hyperparameter tuning in the inner cross-validation loop. For final model comparison, the mean AUROC across the outer folds was calculated.

First, we tested boosting-based models that can handle missing values, including extreme gradient boosting (XGBoost), light gradient boosting machine (LightGBM), and categorical boosting (CatBoost). These models were trained and evaluated using the full set of CGP variants, which allows missing values to remain in the input features.

Second, to enable comparisons with widely used machine-learning methods that do not tolerate missing values, we performed an additional evaluation using a complete-case dataset where variants with missing values were excluded. In this analysis, the above three models were evaluated alongside random forest, k-nearest neighbors, logistic regression, naive Bayes, and support vector machines.

The hyperparameter settings for each model are described below.

**1. Extreme Gradient Boosting (XGBoost)**

XGBoost was applied to both the complete-case dataset and the dataset in which missing values were retained. Models were built using the XGBClassifier from the "xgboost" library. Hyperparameters were optimized through a randomized search, including max_depth (maximum depth of each tree), learning_rate (step size shrinkage to prevent overfitting), n_estimators (number of boosting rounds), "subsample" (fraction of samples used per tree), colsample_bytree (fraction of features used per tree), "gamma" (minimum loss reduction required for further splitting), reg_alpha" (L1 regularization term), and reg_lambda (L2 regularization term).

**2. Light Gradient Boosting Machine (LightGBM)**

LightGBM was applied to both the complete-case dataset and the dataset in which missing values were retained. Models were implemented using the "lightgbm" library. Hyperparameters were optimized through a randomized search, considering the following parameters: "subsample" (fraction of data used for training each iteration), learning_rate (step size for gradient updates), max_depth (maximum depth of individual trees), num_leaves (maximum number of leaves in a tree), colsample_bytree (fraction of features used for training each tree), and n_estimators (number of boosting rounds).

**3. Categorical Boosting (CatBoost)**

CatBoost was applied to both the complete-case dataset and the dataset in which missing values were retained. Models were built using the CatBoostClassifier from the “catboost” library, with log-loss used as the training objective. Hyperparameters were tuned using a randomized search, including “depth” (tree depth controlling model complexity), learning_rate (step size for boosting), “iterations” (number of boosting iterations), “subsample” (fraction of samples used for each tree), “rsm” (fraction of features sampled per tree), min_data_in_leaf (minimum number of samples per leaf), and l2_leaf_reg (L2 regularization strength).

**4. Random Forest**

The random forest model was implemented using the RandomForestClassifier from the sklearn.ensemble module. Hyperparameter tuning was performed via grid search, optimizing n_estimators (number of trees) and max_features (number of features to consider when determining the best split).

**5. k-Nearest Neighbors (k-NN)**

The k-NN model was used with the KNeighborsClassifier from the sklearn.neighbors module. Hyperparameters were tuned through a grid search for n_neighbors (number of neighbors considered).

**6. Logistic Regression**

Logistic regression was implemented using the LogisticRegression class from sklearn.linear_model. Hyperparameters were optimized through a random search for “C” (inverse of regularization strength) and “penalty” (norm used in the penalization).

**7. Naive Bayes**

We applied the naive Bayes algorithm using the scikit-learn library. The Gaussian Naive Bayes variant was used, assuming that the features follow a normal distribution. No hyperparameter tuning was required, as naive Bayes relies on prior probabilities and the likelihood of features given each class.

**8. Support Vector Machine (SVM)**

The SVM model was implemented with the SVC class from sklearn.svm. Hyperparameters such as “C” (regularization parameter), “kernel” (kernel type), and “gamma” (kernel coefficient) were optimized using a randomized search.

**Table S1. Patient Counts by Institution**

| **Institution, *n* (%)** | **Patients (n = 1271)** |
|---|---|
| Institute of Science Tokyo Hospital | 644 (50.1) |
| Tokyo Metropolitan Tama Medical Center | 234 (18.4) |
| Japanese Red Cross Musashino Hospital | 186 (14.6) |
| Tsuchiura Kyodo General Hospital | 123 (9.7) |
| Yokosuka Kyosai Hospital | 84 (6.6) |

#### Table S2. Variant Counts by Annotation Category and Test Type

| Test type | known pathogenic (%) | VUS (%) | Total |
|---|---|---|---|
| **FoundationOne CDx** | 1483 (17.7) | 6898 (82.3) | 8381 |
| **FoundationOne Liquid CDx** | 163 (10.8) | 1344 (89.2) | 1507 |
| **OncoGuide NCC Oncopanel System** | 156 (53.1) | 138 (46.9) | 294 |
| **Guardant360 CDx** | 12 (28) | 31 (72) | 43 |
| **Total** | 1814 (17.7) | 8411 (82.3) | 10225 |

**Abbreviation**: VUS, variants of unknown significance.

#### Table S3. Annotation Categories and Missing Scores of *In Silico* Tool

| Tools | known pathogenic (%) | VUS (%) | Missing (%) |
|---|---|---|---|
| **AlphaMissense** | 1804 (17.6) | 8141 (79.6) | 280 (2.7) |
| **EVE** | 1463 (14.3) | 3788 (37.0) | 4974 (48.6) |
| **ESM1b** | 1811 (17.7) | 8346 (81.6) | 68 (0.6) |
| **PolyPhen-2 HVAR** | 1757 (17.2) | 7883 (77.1) | 585 (5.7) |
| **SIFT** | 1773 (17.3) | 8185 (80.0) | 267 (2.6) |
| **VEST4** | 1770 (17.3) | 8332 (81.5) | 123 (1.2) |

**Note:** “Missing” indicates the number of variants for which a score was not available from each tool.

**Abbreviations**: VUS, variants of unknown significance.

#### Table S4. Two-way ANOVA Results for TMB and Variant Type

| Source | Sum Sq | df | Mean Sq | F | *P* value | $\eta^2p$ |
|---|---|---|---|---|---|---|
| TMB | 7573 | 1 | 7573 | 1097 | < .001 | 0.32 |
| Variant type | 17175 | 1 | 17175 | 2487 | < .001 | 0.52 |
| TMB×Variant type | 4175 | 1 | 4175 | 605 | < .001 | 0.21 |

**Abbreviations**: Sum Sq, sum of squares; df, degrees of freedom; Mean Sq, mean squares; $\eta^2p$, partial eta squared; TMB, tumor mutational burden; ANOVA, analysis of variance.

#### Figure S1. Variant Selection Workflow

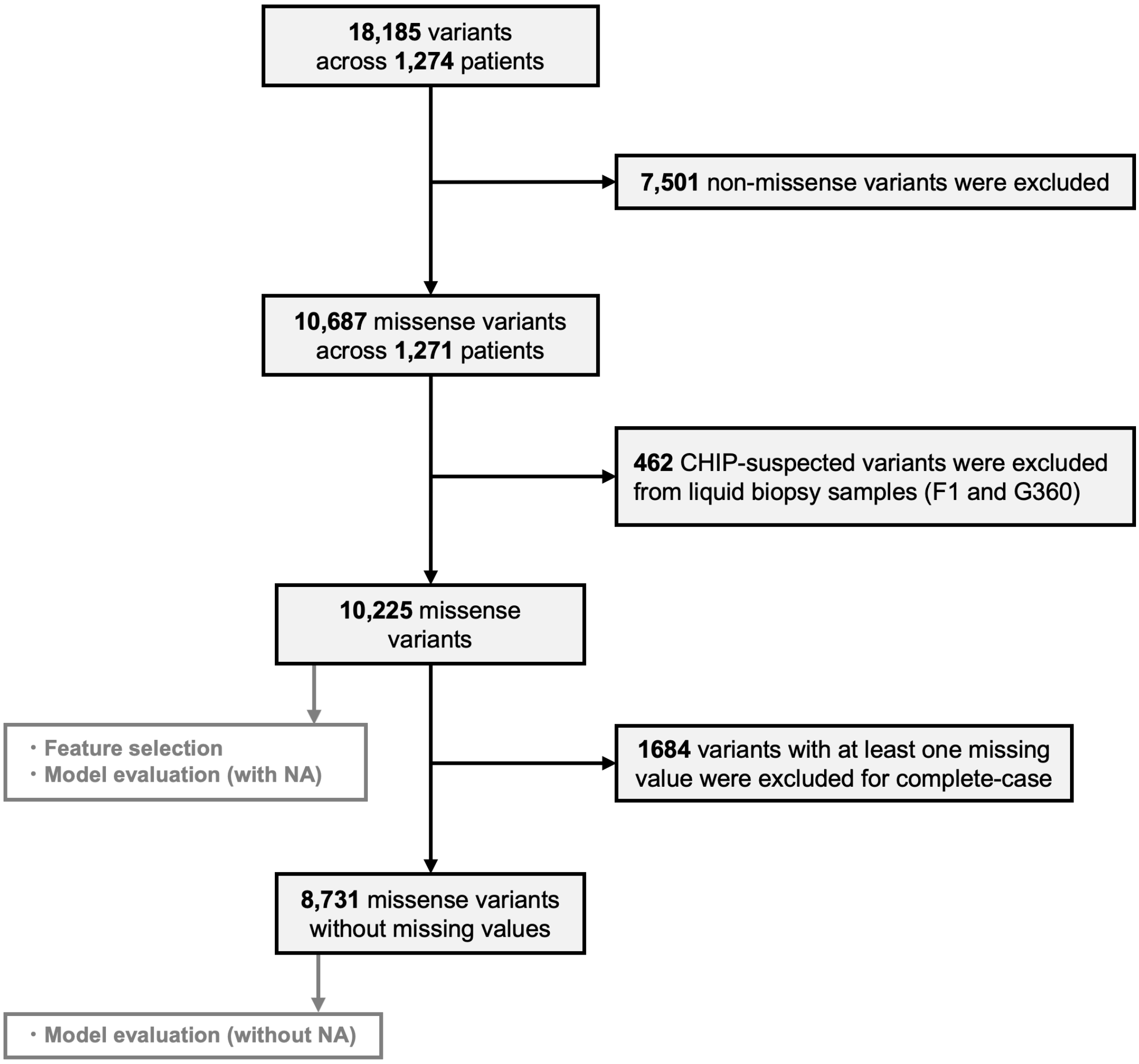


Schematic representation of variant inclusion and exclusion process in this study. The flowchart shows the number of variants analyzed in each section, with inclusion and exclusion criteria.

#### Figure S2. Model Performance on Complete-Case Data

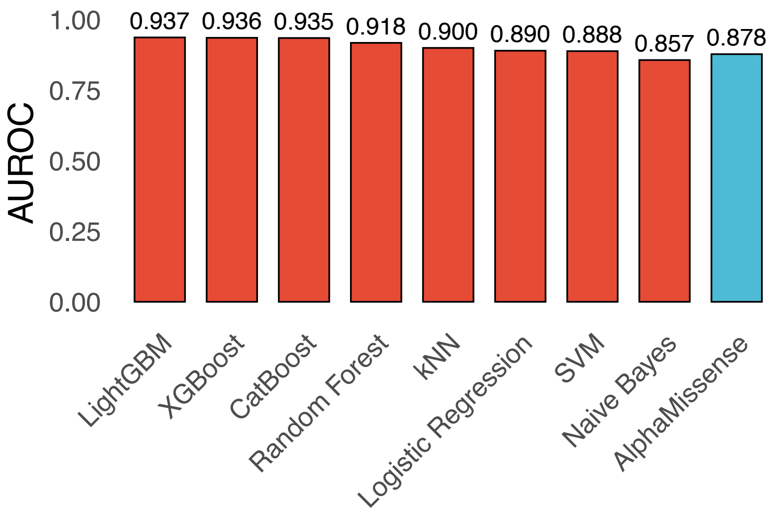


Comparison of the AUROC across multiple machine-learning models evaluated using complete-case data. In addition to the boosting-based models assessed in the main analysis, we included several widely used machine-learning methods that do not tolerate missing values, including random forest, kNN, logistic regression, naive Bayes, and SVM. All models were evaluated on the same dataset restricted to variants without missing values to enable a fair comparison. All boosting-based models consistently demonstrated superior performance compared with any other non-boosting methods, with LightGBM achieving the top performance. Among models that do not handle missing values, random forest achieved the highest AUROC.

**Abbreviations:** AUROC, area under the receiver operating characteristic curve; XGBoost, extreme gradient boosting; LightGBM, light gradient boosting machine; CatBoost, categorical boosting; kNN, k-nearest neighbors, SVM, support vector machines.

**Figure S3. Ablation Study of the Machine Learning Model**

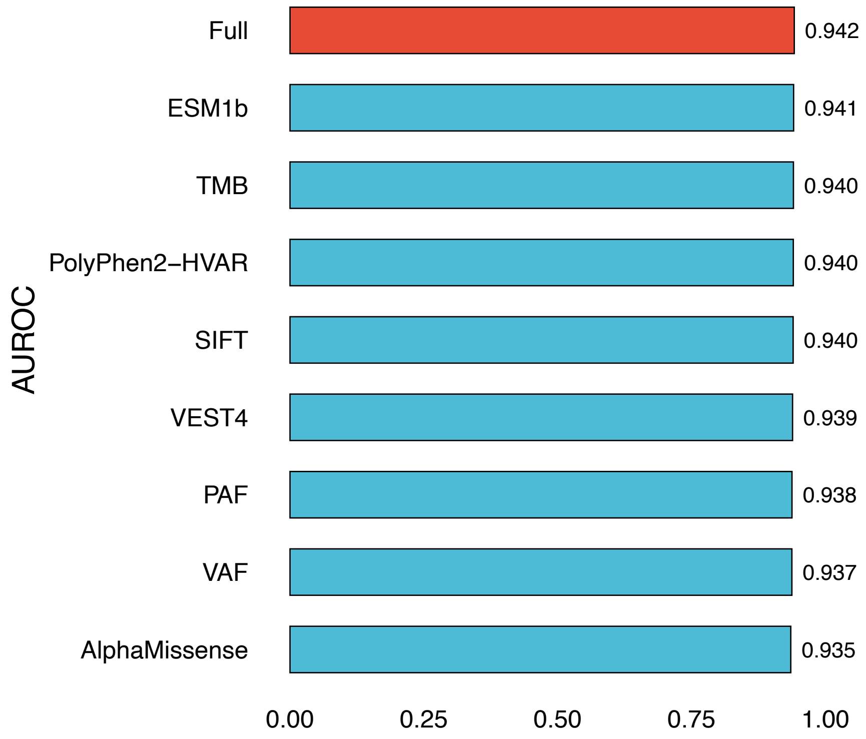


Comparison of the AUROC between the full model (red) and ablated models trained with one feature excluded (blue). The y-axis indicates the excluded feature, and “Full” represents the model trained using all features. The x-axis shows AUROC values. The full model achieved the highest AUROC among all ablated models.

**Abbreviation:** AUROC, area under the receiver operating characteristic curve,

#### Figure S4. Performance of *In Silico* Tools on TCGA Variants Based on ClinVar Labels

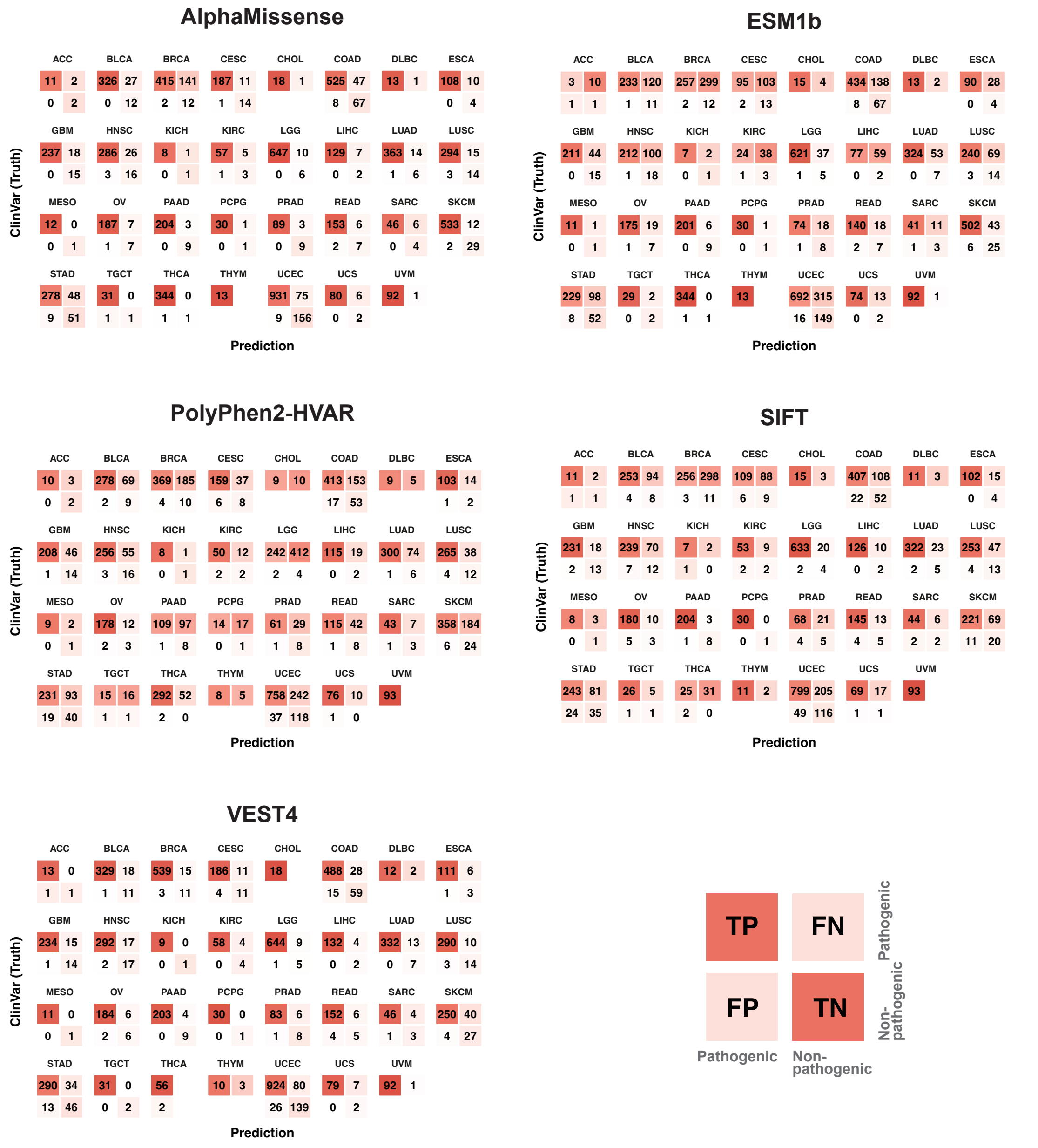


Confusion matrices for AlphaMissense, ESM1b, PolyPhen2-HVAR, SIFT, and VEST4 evaluated on TCGA missense variants with ClinVar annotations classified as pathogenic (“Pathogenic” and “Likely Pathogenic”) or benign (“Benign” and “Likely Benign”). ClinVar classifications were used as ground truth. For each tool, the classification threshold was determined using the Youden index based on the CGP dataset.

**Abbreviations:** TCGA, The Cancer Genome Atlas; CGP, comprehensive genomic profiling; TP, true positive; FN, false negative; FP, false positive; TN, true negative.

#### Figure S5. Comparison of MARiO and a Model Without Adjusted TMB using TCGA Variants Based on ClinVar Labels

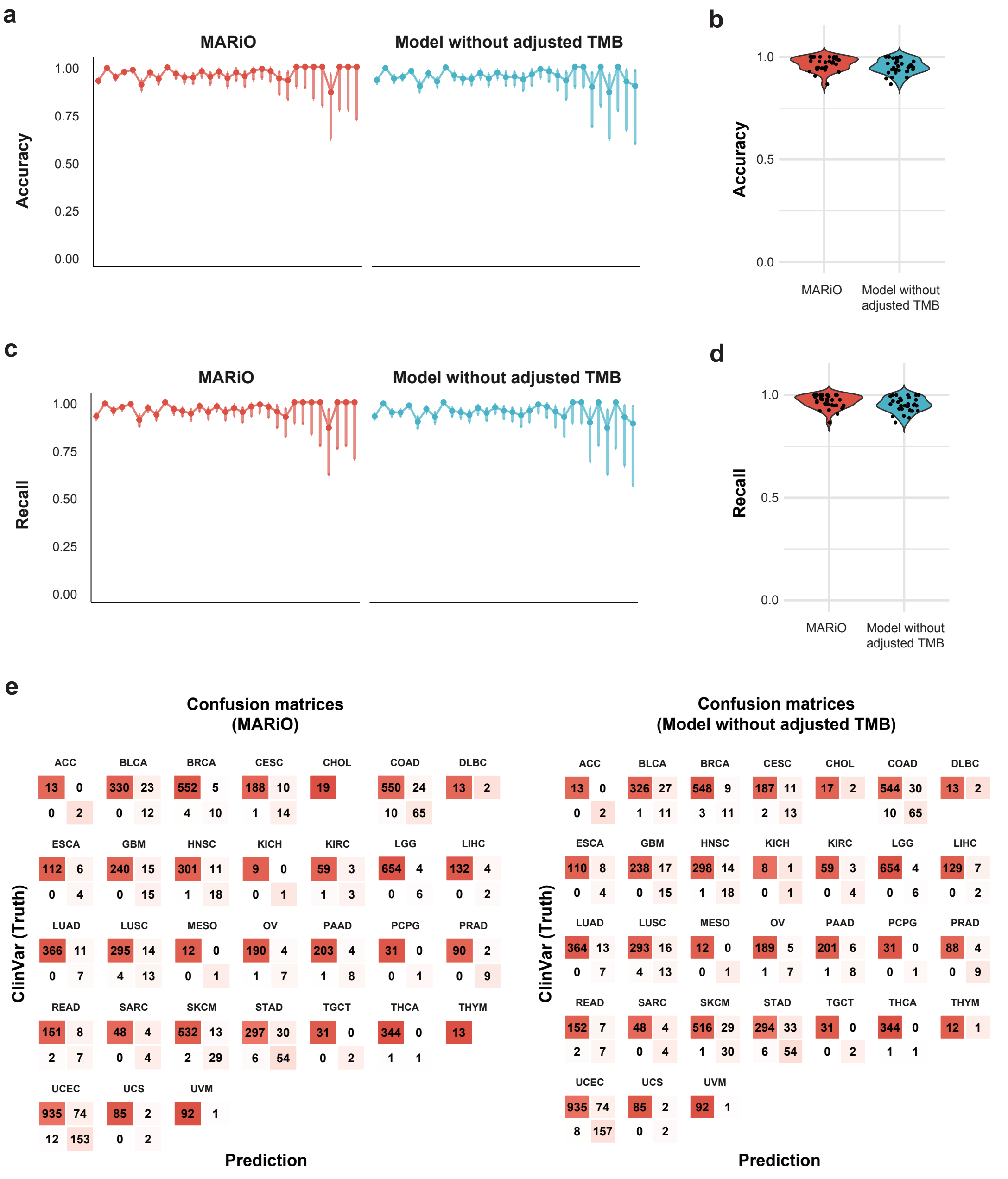


Performance was evaluated on TCGA missense variants with ClinVar annotations classified as pathogenic or benign.

**a** Line plots showing accuracy across cancer types. Each point represents the performance metric for each cancer type, with vertical bars indicating 95% confidence intervals calculated using the Wilson's score method. Cancer types are ordered from left to right by decreasing sample size, as follows: UCEC, LGG, COAD, SKCM, BRCA, STAD, LUAD, BLCA, THCA, HNSC, LUSC, GBM, PAAD, CESC, OV, READ, LIHC, ESCA, PRAD, UVM, UCS, KIRC, SARC, TGCT, PCPG, CHOL, ACC, DLBC, MESO, THYM, KICH.
**b** Violin plots showing the distribution of accuracy.
**c** Line plots showing recall across cancer types, with the same format and cohort ordering as in panel a.
**d** Violin plots showing the distribution of recall.
**e** Confusion matrices for MARiO and the model without adjusted TMB.

**Abbreviations:** MARiO, Missense Alteration Risk for Oncogenicity; TMB, tumor mutational burden; TCGA, The Cancer Genome Atlas.

### Figure S6. Validation of MARiO Scores Using the Cancer Hotspots Catalog

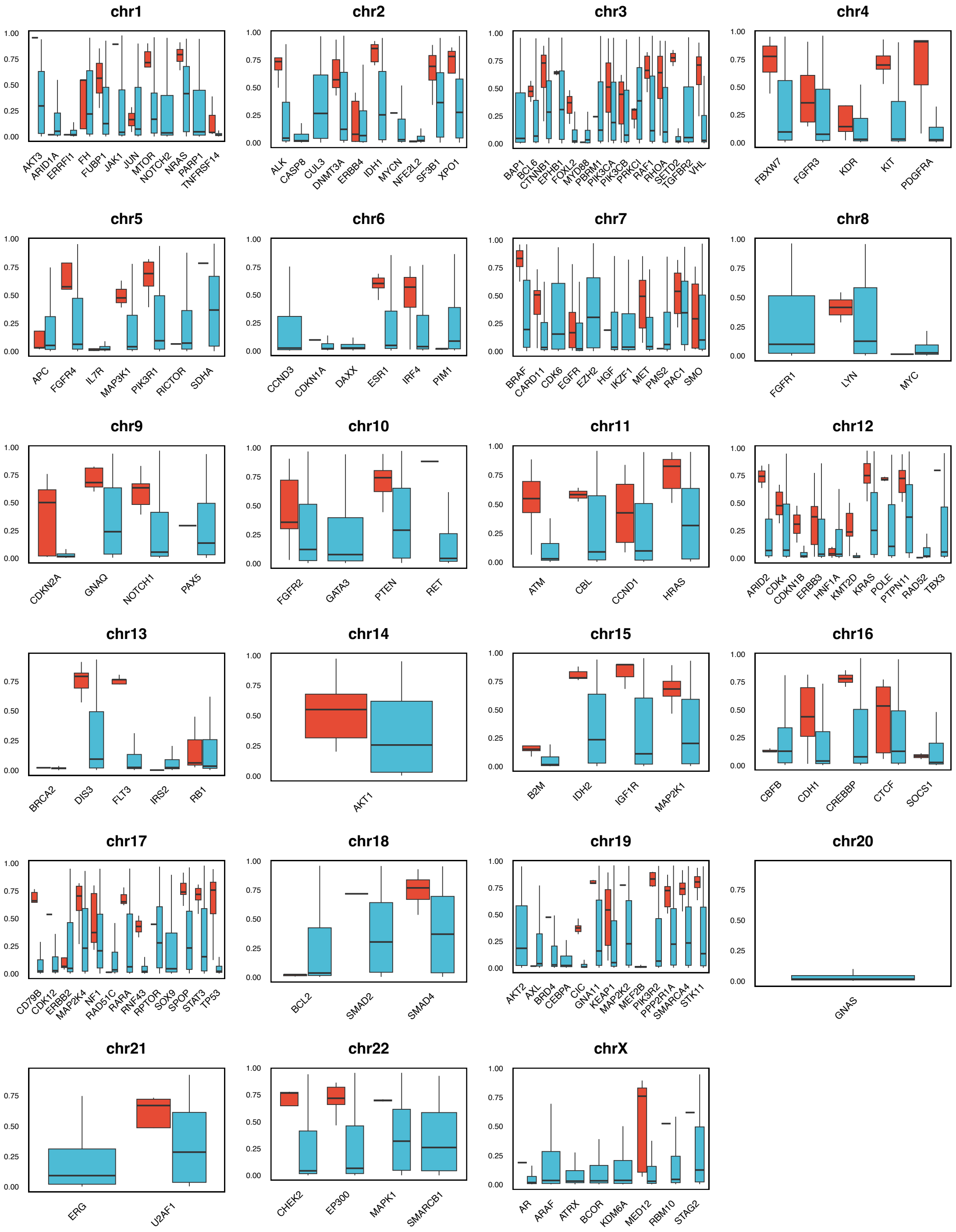


Distribution of MARiO scores for hotspot and non-hotspot variants across genes. Each panel corresponds to a chromosome, with genes shown on the x-axis and MARiO scores on the y-axis. Hotspot annotations were obtained from the Cancer Hotspots catalog. Red indicates hotspot variants, and blue indicates non-hotspot variants.

#### Figure S7. Cohort for Treatment-Related Analyses

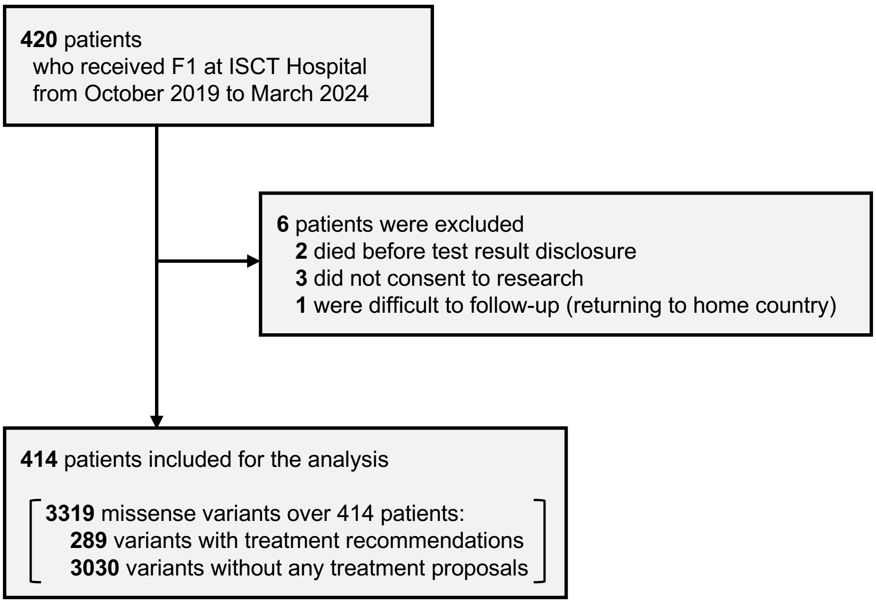


A total of 420 patients received CGP testing (F1) at ISCT Hospital from October 2019 to April 2024. Six patients were excluded based on predefined exclusion criteria, resulting in 414 patients included in the treatment-related information analysis.

**Abbreviations:** ISCT, Institute of Science Tokyo; CGP, comprehensive genomic profiling; F1, FoundationOne CDx.
